# Building a genome-based understanding of bacterial pH preferences

**DOI:** 10.1101/2023.01.24.524446

**Authors:** Josep Ramoneda, Elias Stallard-Olivera, Michael Hoffert, Claire C. Winfrey, Masumi Stadler, Juan Pablo Niño-García, Noah Fierer

## Abstract

The environmental preferences of many microbes remain undetermined. This is the case for bacterial pH preferences, which can be difficult to predict *a priori* despite the importance of pH as a factor structuring bacterial communities in many systems. We compiled data on bacterial distributions from five datasets spanning pH gradients in soil and freshwater systems (1470 samples in total), quantified the pH preferences of bacterial taxa across these datasets, and compiled genomic data from representative bacterial taxa. While taxonomic and phylogenetic information were generally poor predictors of bacterial pH preferences, we identified genes consistently associated with pH preference across environments. We then developed and validated a machine learning model to estimate bacterial pH preferences from genomic information alone, a model which could aid in the selection of microbial inoculants, improve species distribution models, or help design effective cultivation strategies. More generally, we demonstrate the value of combining biogeographic and genomic data to infer and predict the environmental preferences of diverse bacterial taxa.

## Introduction

Predicting the environmental preferences of organisms is an important goal in ecology. If we know the conditions under which a given taxon can thrive, we can better predict biogeographical distributions (*1*), guide ecological restoration efforts (*2*), design effective probiotics (*3*), and understand taxon-specific responses to global change factors (*4*). Unfortunately, the environmental preferences of most microbial taxa, and the genomic attributes associated with those preferences, often remain undetermined (*5*)–(*6*). One reason for this is that most microorganisms, particularly those found in non-host associated environments, can be difficult to cultivate *in vitro* (*7*), making it difficult to measure environmental preferences directly. Even for those taxa that can be cultivated, quantifying how microbial growth rates vary across broad environmental gradients can be time consuming and the environmental gradients created *in vitro* may not necessarily mimic those found *in situ*. However, when direct information on environmental preferences can be collected, such information can be very useful for predicting microbial distributions and functions across space and time (e.g. (*8*–*11*)).

Even without the direct measurement of environmental preferences, it is feasible to infer some environmental preferences from genomic information (*12*). We can leverage the information contained in curated genomic databases, which can include both cultivated and uncultivated microbial taxa (*13*), to infer the specific environmental preferences of uncharacterized microorganisms (*14*). For example, genomic information from isolates whose environmental preferences have been measured *in vitro* has been used to infer the preferences of bacteria across gradients in oxygen (*15*) and temperature (*12*). Using genomic information to determine the environmental preferences of microbial taxa can have important ramifications. For example, we could improve our ability to predict community assembly across different environmental gradients, identify the conditions under which specific taxa can thrive, and better optimize media conditions to improve the cultivability of fastidious taxa. However, genome-based inferences of environmental preferences can be difficult to validate (especially for uncultivated taxa) and we do not always know which genes or other genomic attributes are associated with adaptations to specific environmental conditions of interest.

Consider microbial preferences for specific pH conditions. To our knowledge, it is not currently possible to predict bacterial pH preferences from genomic information alone, although we know that pH is often a key factor determining the niche space occupied by microorganisms. The distributions of specific bacterial taxa, and the overall composition of bacterial communities, are often strongly associated with gradients in pH, as has been observed in a wide range of environments including soils (*16*–*19*), freshwater (*20*–*22*), and geothermal systems (*23*). Despite the importance of this environmental factor, the pH preferences of most bacterial taxa remain undetermined, although we know that bacterial pH preferences can vary widely (*24*). This knowledge gap is particularly evident in non-host associated systems which are often dominated by taxa that are difficult to cultivate and study *in vitro*. Even across cultivated taxa, pH tolerances and pH optima for growth are rarely determined experimentally and most cultivation media likely select for taxa that grow at near-neutral pH conditions (*25*). We note that ‘pH preference’ across a given gradient is akin to the ‘realized niche’ of a population (as opposed to the ‘fundamental niche’), where pH preference is the pH at which an organism achieves maximal relative abundances in nature (*17*). This relative abundance is determined by the metabolically optimal pH for growth as well as other biotic or abiotic constraints. For example, a given taxon may grow optimally at around pH 7 under controlled conditions, but its pH preference could be lower if its abundance in a given environment is maximized at pH 6 due to its interplay with other factors.

In the absence of whole genome sequence data, one might still be able to predict bacterial pH preferences from taxonomic and phylogenetic information alone. Shifts in bacterial community composition across pH gradients can be evident at broad levels of taxonomic resolution (*26*, *27*), suggesting some degree of conservatism in the traits related to pH adaptation. However, closely related taxa can have very distinct pH preferences (*28*), such as bacteria within the phylum Acidobacteria which tend to be more abundant in lower pH soils (*27*), a pattern that does not necessarily hold for sub-groups within the phylum (*29*, *30*). Thus, it is not clear if taxonomy and phylogenetic information are sufficient to predict bacterial pH preferences. Previous studies have focused on identifying common adaptations to changes in pH conditions across different taxonomic groups, and the genes or transcripts associated with those adaptations (*28*, *31*–*33*). This generates the expectation that the presence or absence of specific functional genes can be used to predict bacterial pH preferences. An approach that integrates biogeographic distributions of multiple individual taxa with genomic information across environmental gradients can address whether taxonomic, phylogenetic, or genomic information can be used to predict microbial environmental preferences.

We set out to determine if bacterial pH preferences are predictable. In other words, we asked whether we could use taxonomic, phylogenetic, or genomic information to predict where along a pH gradient a taxon is most likely to achieve its highest relative abundance (which we define here as its ‘pH preference’), searching for patterns that are generalizable across distinct environment types. To do so, we used information on bacterial distributions across 5 independent datasets that span large pH gradients in soil and freshwater systems to infer the putative pH preferences of the bacterial taxa found in these environments. We then used this information to assess the degree to which bacterial pH preferences are phylogenetically and taxonomically conserved. To determine the genomic features associated with adaptations to pH, we analyzed representative genomes from taxa with varying pH preferences, as inferred from our cultivation-independent analyses, and identified genes that are consistently associated with differences in bacterial pH preference across environments. Finally, we developed and validated a machine learning model that enables the accurate identification of bacterial pH preferences from the presence or absence of 56 functional genes, making it feasible to infer pH preferences for both uncultivated and cultivated taxa. More generally, we demonstrate how our workflow (Fig. 1) can be used to investigate other bacterial environmental preferences, and the genes associated with environmental adaptations, while overcoming the limitations of cultivation-based experimental approaches to expand our trait-based understanding of microorganisms.

**Fig. 1.**
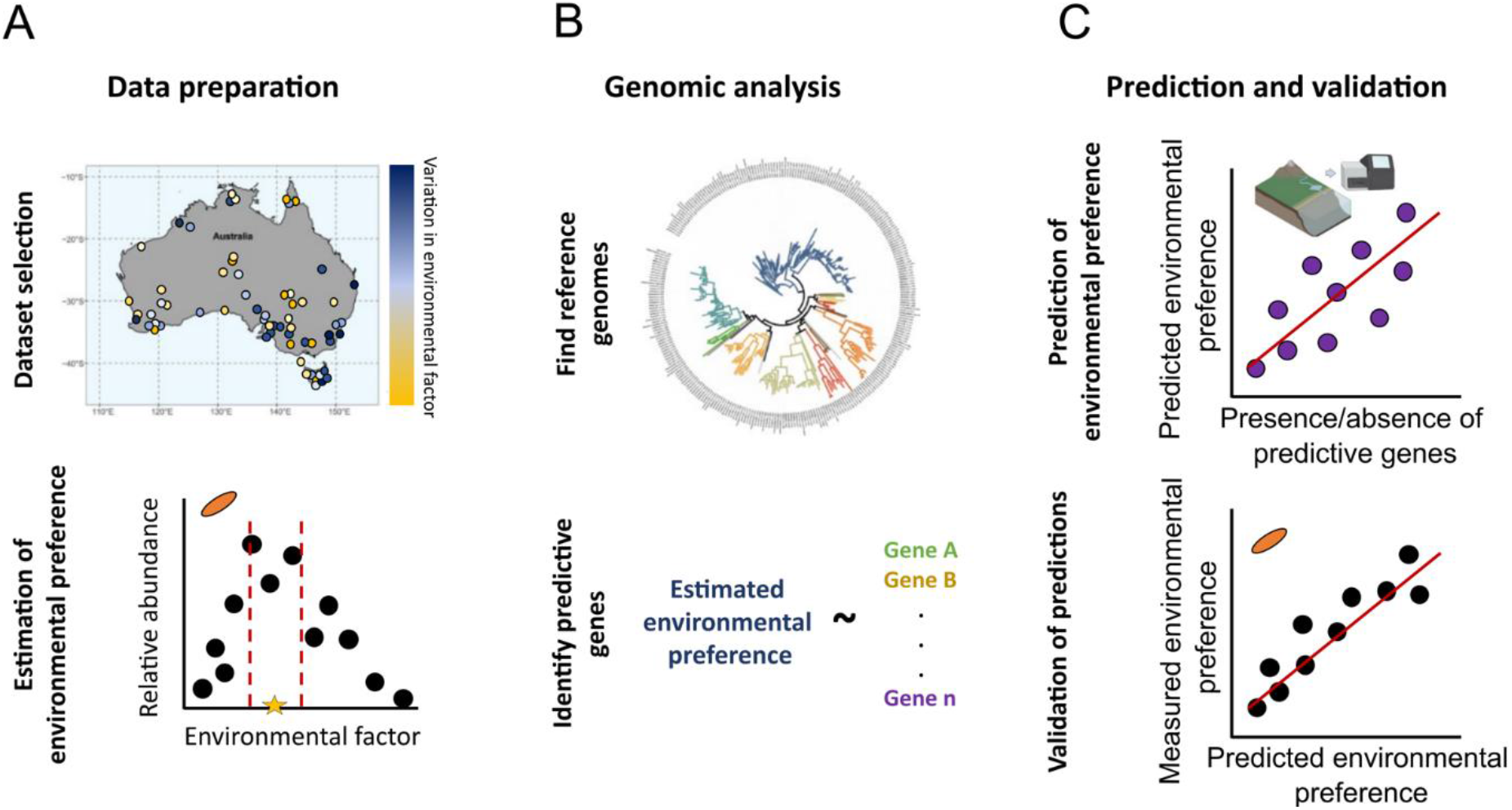
Workflow to predict environmental preferences from genomic and biogeographical information. (A) Marker gene sequencing datasets that include samples spanning a broad range in the environmental factor of interest (pH in our case) are chosen and rare taxa are excluded based on occurrence. The remaining taxa are then used for the estimation of the environmental conditions at which the relative abundances of individual taxa are highest using either bootstrapping or generalized additive model (GAM) approaches. (B) The taxa for which an environmental preference has been statistically estimated are then queried against curated genome databases. Genomes matching the query sequence at high similarity (e.g. at 99.6% in this study) are considered representative of the taxa. (C) All genes present in those genomes are then tested for their relationship to environmental preference, and those that are significantly associated with environmental preference (e.g. sharing the same statistically significant positive or negative relationship with pH across datasets) are incorporated into a machine learning modelling framework. Predictions can be validated using an independent test set of the data, from pH preferences estimated in independent datasets, or from phenotypic assays.

## Results and Discussion

### Overview of the approach

We first inferred bacterial pH preferences using biogeographical information from natural pH gradients found in distinct systems (soil and freshwater environments) (Fig. 1A). We acknowledge that pH is unlikely to be the only factor influencing the distributions of bacteria in these systems and that the pH preferences of many taxa could not be inferred using this approach as there are other biotic or abiotic factors that are of similar, or greater, importance in shaping their distributions. However, we note that this approach is similar to the approach routinely used in plant and animal systems to quantify the relationships between environmental factors and growth optima or tolerances (*39*, *55*, *56*). After inferring pH preferences from the biogeographical information, we then matched the 16S rRNA gene sequences from those taxa for which pH preferences could be inferred to representative genomes, identifying sets of genes that are consistently associated with pH preference based on their presence or absence in a given genome (Fig. 1B). These genes were then incorporated into a machine learning modelling framework (Fig. 1C) to predict pH preferences, with the approach validated using independent test sets.

The biogeographical analyses included 16S rRNA gene sequence data from a total of 795 soil samples and 675 freshwater samples spanning a pH range from 3 to 10 with a total of 250,275 ASVs (amplicon sequence variants) included in downstream analyses (Fig. S1; Suppl. Table 1). These data came from 5 independent datasets, and in all cases, pH was an important driver of overall bacterial community composition, with Mantel rho values ranging from 0.37 (ROMAINE, freshwater) to 0.78 (PAN, tropical forest soils) (Fig. S1). These patterns are not surprising considering that pH is often observed to have a strong influence on bacterial community structure in many systems (*27*, *37*) and considering that each of these sample sets were specifically selected to span broad gradients in pH. We also note that a broad diversity of bacterial taxa was found within and across each of the 5 sample sets (Fig. S2A), which is important as we were trying to identify patterns in pH preferences across a broad array of taxa. Using our conservative approach, we were able to estimate the pH preferences of bacterial taxa in these environmental samples for 0.5-4.9% of all ASVs per dataset (Table S1). The analyses of the taxonomic and phylogenetic signals in bacterial pH preferences and the search for representative genomes from these taxa was ultimately based on a total of 4568 ASVs (468 – 1614 ASVs per dataset) spanning 38 bacterial phyla.

### Are taxonomic and phylogenetic information good predictors of the pH preference of bacterial taxa?

Taxonomy was a poor predictor of bacterial pH preferences (Fig. 2). In almost every phylum there were ASVs with very distinct pH preferences and there were few cases where a high proportion of ASVs from a particular phylum were found to have a similar pH preference. For example, in the ROMAINE freshwater dataset, many ASVs assigned to the phylum Acidobacteria did exhibit a general preference for acidic pH conditions, but this observation was not consistent across datasets (Fig. 2). Thus phylum-level information provides relatively little insight into bacterial pH preferences, most likely because taxa within a given phylum can have divergent pH preferences (as noted previously for different acidobacterial subdivisions (*30*, *38*)). We observed similar patterns at finer levels of taxonomic resolution. For example, ASVs within some of the most ubiquitous bacterial families observed across these 5 datasets (Xanthobacteraceae in soil and Chitinophagaceae in freshwater systems) included ASVs with inferred pH preferences ranging from 4.01 to 8.20 and from 4.63 to 8.35, respectively.

**Fig. 2.**
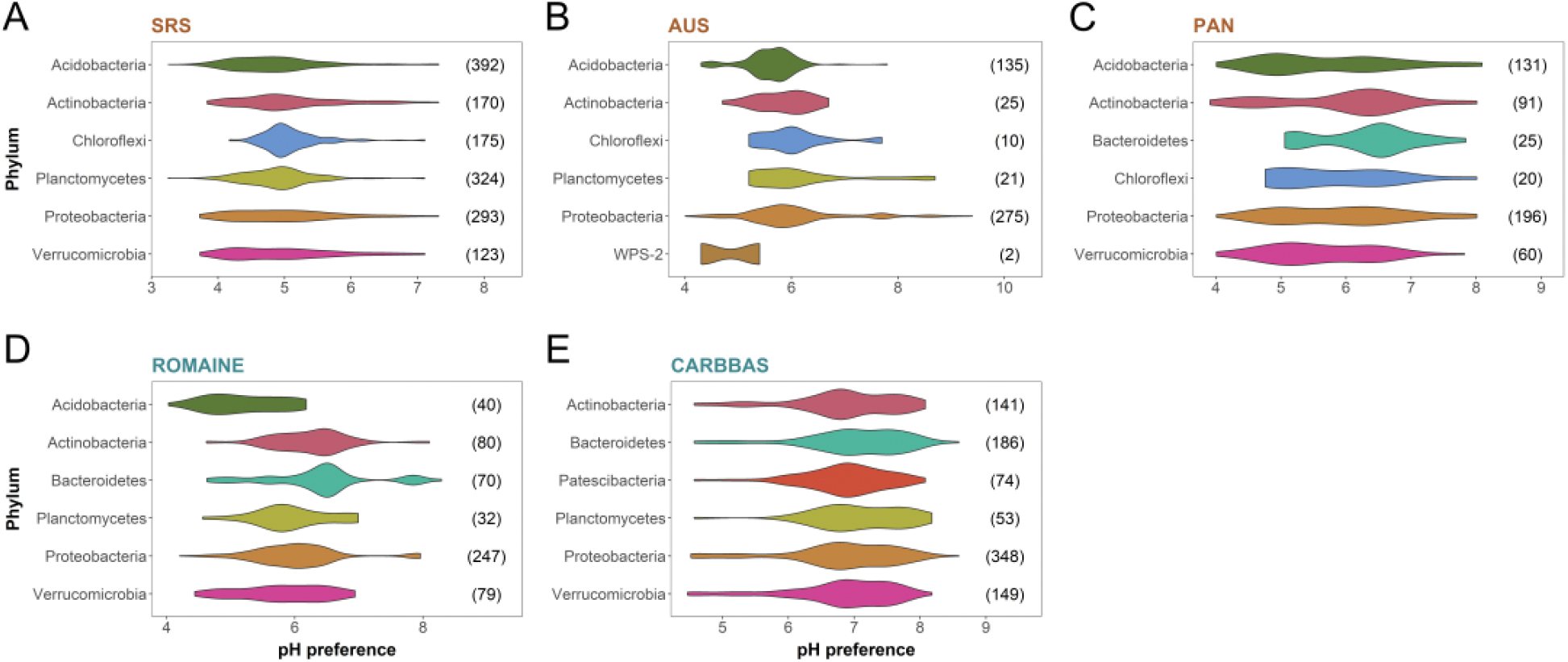
Range of pH preferences of those ASVs from which pH preferences could effectively be inferred from the biogeographical patterns in the marker gene sequencing data. The taxonomic assignments of the ASVs are shown at the phylum level with more complete taxonomic information included in Figure S2A. Values at the end of the boxplots correspond to the number of ASVs with estimated pH preferences in each phylum. The length and width of the violins depict the entire range of pH preferences and the most frequent pH preferences of taxa within each phylum, respectively.

Consistent with the taxonomy-based results, we also found that bacterial pH preferences were not readily predictable from phylogenetic information. In other words, there was minimal phylogenetic conservation in pH preferences. Although we detected a significant phylogenetic signal in pH preferences for all but one dataset (Fig. 3, Fig. S3), the signal was relatively weak (Pagel’s λ 0.22-0.78) (Fig. S3). This is qualitatively evident from the phylogenetic trees associated with each dataset which show that even closely related taxa often had very distinct pH preferences. For example, within the Proteobacteria phylum some clusters of ASVs with similar preference for higher or lower pH conditions were indeed observed in the phylogenies from both AUS (soil) and ROMAINE (freshwater) datasets, but these clusters were not evident in the other datasets (Fig. 3). These observations were further confirmed by the phylogenetic correlogram analyses which show that the Moran’s I autocorrelation values were low in all cases (<0.25 (*39*)), even at shallow phylogenetic depth (Fig. S2B; Fig. S3). Our results are in line with previous work that found no phylogenetic conservation of pH preferences across both cultured bacteria (*14*) and uncultured bacteria (*40*). Neither taxonomic nor phylogenetic information are generally useful for determining pH preferences without additional information, highlighting the potential value of incorporating functional gene information to better predict bacterial pH preferences.

**Fig. 3.**
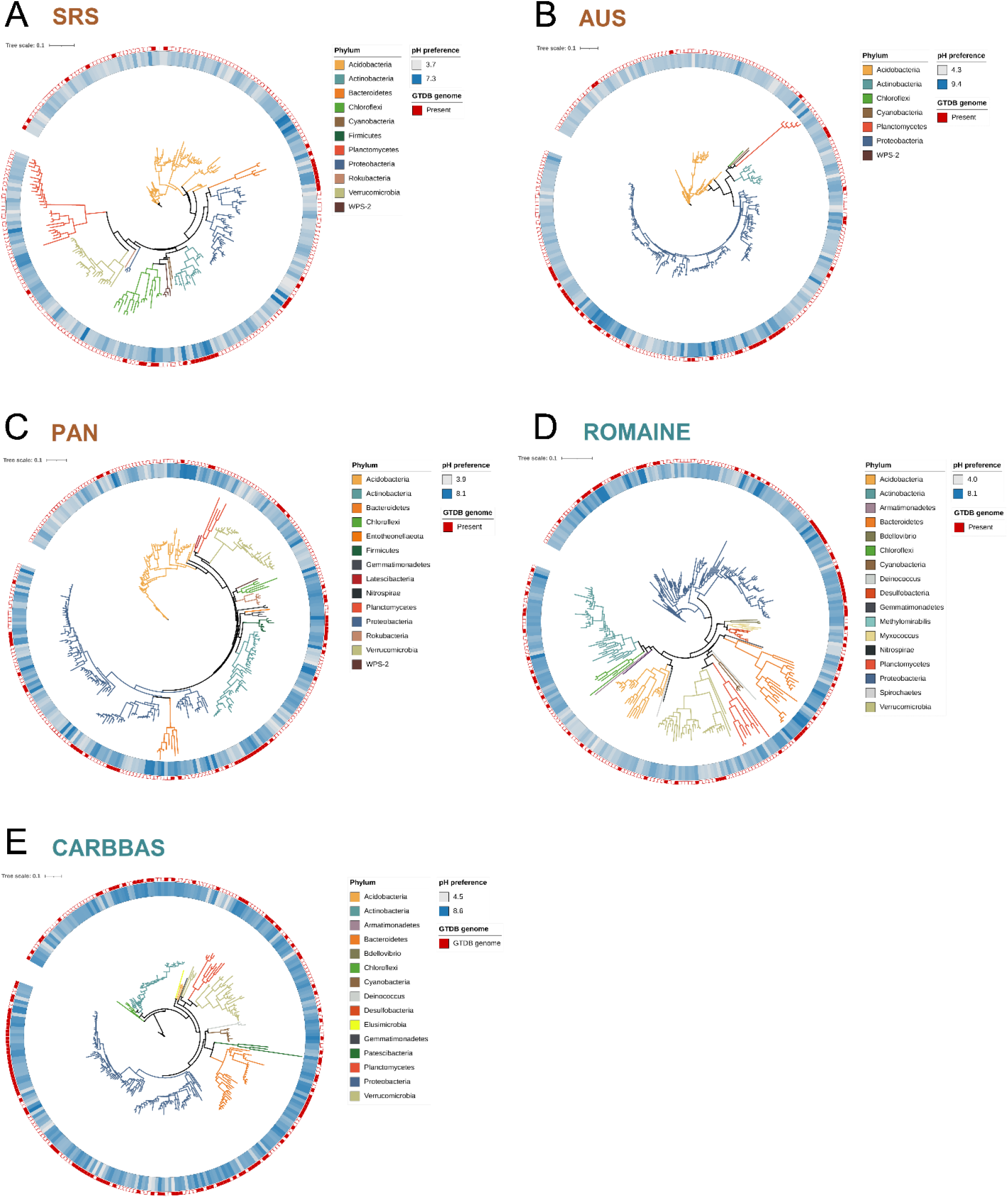
Phylogenetic distributions of the most abundant amplicon sequence variants (ASVs) with estimated pH preferences in each dataset. Clades are colored by the affiliation at the phylum level. The inner ring depicts the estimated pH preference of the ASVs, and the outer ring indicates whether a genome with no more than one nucleotide mismatch to the amplicon sequence is available in the Genome Taxonomy Database (GTDB). The displays are maximum likelihood phylogenetic trees, and these trees only show the 300 most abundant ASVs per dataset, with the complete set of ASVs (numbers indicated in Figure 3) used for the calculation of the phylogenetic signal in pH preferences (see text and Figure S3). The minimum pH preference is depicted in white, and the maximum is depicted in blue according to the legend.

### Associations between bacterial pH preferences and functional genes

We next analyzed genomes representative of those ASVs with inferred pH preferences. Not surprisingly, given the well-recognized biases in reference genome databases (*41*), only a relatively small fraction (6 - 24%) of those ASVs identified in our sample sets had representative genomes in GTDB (*42*) and a significant portion (~35%) of those genomes were metagenome-assembled genomes (MAGs) and single cell-assembled genomes (SAGs) from uncultured taxa (Fig. S4). The number of genome matches obtained ranged from 57 genomes in the AUS soil dataset to 293 genomes in the CARBBAS freshwater dataset (with a total across all 5 datasets of 669 ASV-genome matches representing 580 unique genomes with inferred pH preferences; Table S1, Fig. S5A). From the taxa with inferred pH preferences, the proportion of ASVs with available genome representatives was 10.6% in soils and 22.4%in freshwater on average (Fig. S5A). Still, the taxonomic composition of the available genomes spanned 20 different phyla, with a general predominance of Actinobacteria and Proteobacteria among the matching genomes (Fig. S5B), a result that is unsurprising given that these two phyla are relatively well-represented in genome databases (*5*).

We identified 332 gene types that had the same significant association with inferred pH preference in at least two datasets, while 56 of those genes had the same association to pH in at least three datasets across soil and freshwater habitats (Fig. 4; Table S2). These gene types encoded for proteins known for their involvement in pH tolerance such as ATPases, anion and cation transporters and antiporters, and alkaline and acidic phosphatases (Fig. 4). Generally, bacteria need to maintain pH homeostasis to maintain enzymatic function and cytoplasmic membrane stability, and they generally use four main mechanisms to cope with acid stress (*43*, *44*). Across those genes that we identified as being associated with pH preferences (Fig. 4), we see evidence for all four mechanisms. First, proton-consuming reactions, notably decarboxylation and deamination of amino acids, buffer proton concentrations in the cytoplasm by incorporation of H^+^ to metabolic by-products (*45*, *46*). We identified genes for decarboxylases (AAL_decarboxy, soils), amino acid transporters (AA_permease, freshwater), carboxylate transporters (*TctA*, soils and freshwater), and amino acid deaminases (Queuosine_synth, soils and freshwater) associated with inferred pH preferences. Second, cells will produce basic compounds, such as ammonia released from urea to counter acidity. We found genes assigned to urea membrane transporters (ureide_permeases) overrepresented in taxa with low pH preference in soil and freshwater, as well as a gene for urease (UreE_C, soils) that hydrolyzes urea into ammonia (*47*). Third, bacteria can actively efflux protons to maintain intracellular pH levels. We identified genes for a wide range of cation and anion efflux pumps such as the *Kdp* K^+^ membrane transporters (KdpACD) that were overrepresented in taxa with low pH preference in all habitats. In contrast, Na^+^/H^+^ antiporters (*PhaGF*, *MnhG*, *MrpF*, *YufB*; (*33*)) and anion transporters such as citrate (CitMHS, soils and freshwater) and lactate permeases (freshwater) were overrepresented in taxa with preferences for higher pH. Finally, bacteria also modify the permeability of the cytoplasmic membrane and control the maturation and folding of proteins to limit acid stress. We identified genes for multiple hydrogenase quality control proteins (*HypCD*, *HycI*, *HupF*) as overrepresented in taxa with low pH preference across soils and freshwater, with these genes known to be involved in acid stress response (*47*, *48*). We also identified genes for process-specific proteins that act in a pH-dependent manner such as acidic phosphatases (Phosphoesterase and CpsB_CapC in soils and freshwater; (*32*)). Malik et al., (*28*), who gathered genomic information from soil bacterial communities across a pH gradient of 4-8.5, found very similar genes and related functions to our gene types (summarized in Table S2). Overall, we found evidence for the involvement in bacterial pH tolerance in 30 out of those 56 gene types that were consistently associated with pH preference across habitats (Fig. 4; Table S2). While we cannot confirm that these genes we identified as being associated with pH preferences across multiple datasets actually represent specific bacterial adaptations to pH, our results make it possible to generate new hypotheses about novel genes involved in bacterial responses to pH. Our results expand on previous work by identifying multiple gene functions associated with pH preferences beyond transporter genes (*14*), while these results also confirm well-known associations between functional genes and pH preference in bacteria. Also, since most of our current knowledge regarding the specific genes involved in pH tolerance are derived from *in vitro* studies focused on bacterial pathogens (*49*), our findings provide a basis to extend the investigation of bacterial adaptations to pH more broadly beyond those selected taxa.

**Fig. 4.**
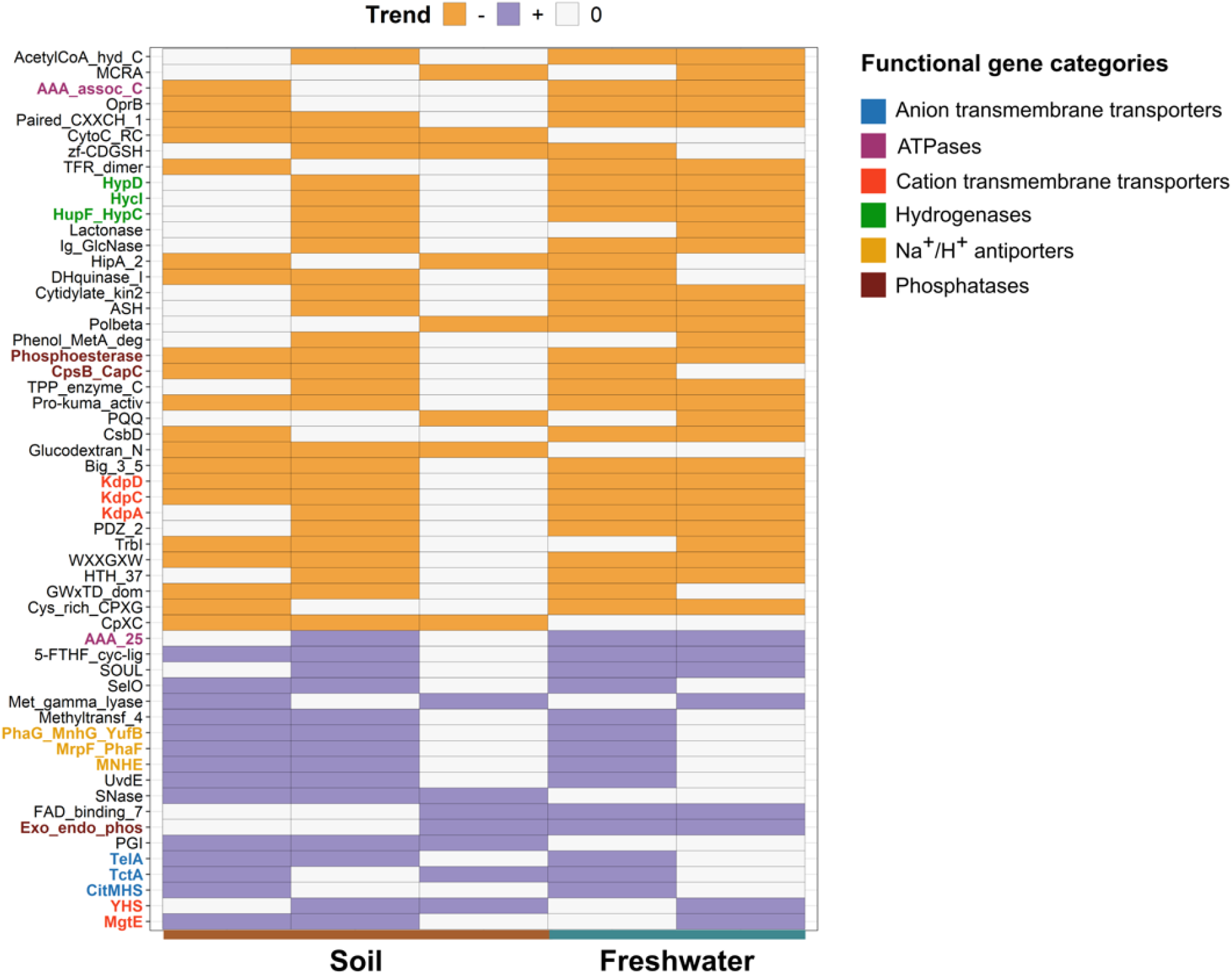
Associations between gene categories and estimated pH preferences in bacteria across soils and freshwater systems. Positive trends (blue cells) indicate that the presence of the gene is associated with inferred bacterial preferences for higher pH, and negative trends (orange cells) indicate that presence of the gene is associated with bacterial preferences for lower pH conditions. We only included gene types that had consistent relationships to pH preference in at least three datasets and/or in both soil and freshwater. We highlighted categories with well-known involvement in pH tolerance and response in bacteria. For a complete summary of the putative functionality, Pfam identifier, and available references linking these specific gene types to pH adaptations see Table S2. The x-axis was sorted left to right in the order of SRS, AUS, PAN, ROMAINE, and CARBBAS datasets, respectively.

### Prediction of bacterial pH preference from genomic information alone

We incorporated the information on presence/absence of the 56 gene types consistently associated with pH preference across habitats into a machine learning modelling framework to predict bacterial pH preferences. Across datasets, we obtained an average R^2^ value of 0.80 for the linear regression between predicted and observed pH preferences using the training data (Fig. 5). These genes encoded very diverse functions, from well-established ones such as transmembrane anion and cation transport, ATPase, and phosphatase activity, to functions less known to be involved in bacterial pH responses such as nucleases for DNA repair, endolysins, or the type V secretion system (Table S2). With only presence/absence information of these 56 genes, we were able to predict the pH preferences of bacterial taxa that had not been used for model training. The validations conducted on a randomly selected subset of the genomes in each dataset (10% of genomes) had a mean absolute error (MAE) of 0.63 (MAE = 0.43 using the training data; Fig. 5), indicating that with available presence/absence information of these 56 genes, this machine learning model can predict the pH preference of a given bacterial taxon with an accuracy of 0.63 pH units. Considering bacterial taxa generally have pH optima within 1 pH unit (*50*, *51*), this error is relatively small. Likewise, the average R^2^ of the linear regression between the observed and predicted pH preferences in the independent validation set was 0.55. Note that, with this model, the absence of a gene is as informative for predictive purposes as gene presence (Fig. S6), and that, due to the lack of data for taxa with estimated pH preferences below pH 4 and above pH 9, the model can only predict accurately within that range. We also note that our model predicted bacterial pH preferences across all 5 datasets, not necessarily pH optima for growth. Observed pH preferences correspond to the pH at which taxa achieve maximal abundance in nature, reflecting both the pH optimum for growth and the biotic and abiotic factors that may constrain bacterial abundances – the ‘realized niche’. We have made a detailed description of this model and how it can be applied to any genome of interest in https://github.com/fiererlab/ph_preference.

**Fig. 5.**
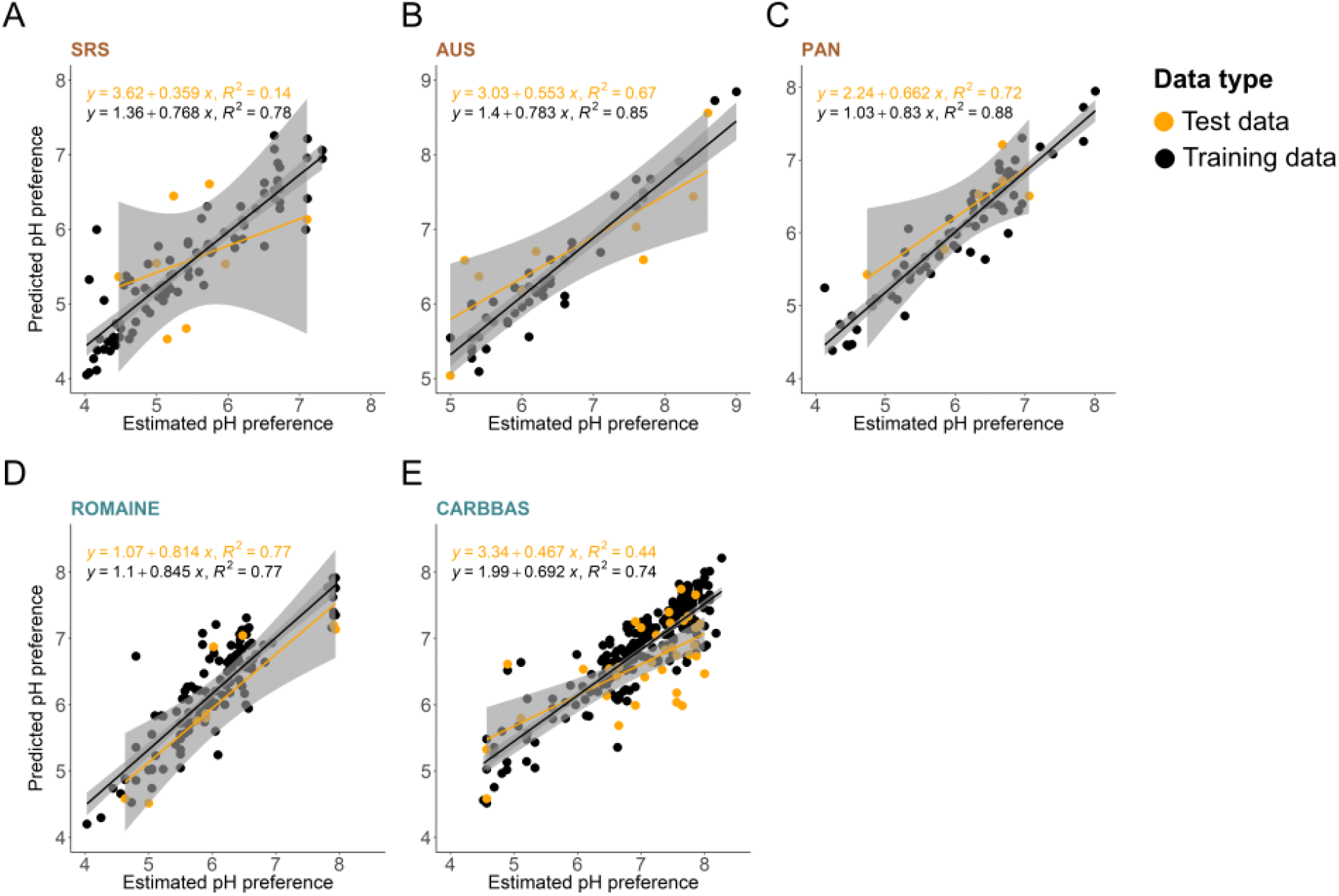
Correlations between the estimated and predicted pH preferences of bacterial taxa based on a gradient boosted decision tree model built on the presence/absence of 56 top predictor gene types. Each panel corresponds to a distinct dataset from either soil (brown) or freshwater (blue). The regressions on the plot include the predictions using the full dataset (black) and the independent validation using 10% of the genomes (orange) not included in the model development. The grey shaded area depicts the standard error around the fitted line.

We further validated our model using information on bacterial pH preferences estimated in a completely independent study of bacterial distributions in soils across the UK (*40*). The linear regression between the statistically estimated pH preference of taxa in the dataset and our model predictions had R^2^ = 0.21 and MAE = 0.93 (Fig. S7). Note that the estimated pH preferences in that study were obtained using a different approach from our study, and that the pH preferences for many taxa were inferred using models with relatively weak fits, a point the authors were careful to acknowledge (*40*). Despite the limitations of this independent dataset and the limitations associated with our model, the correspondence between predicted and observed pH preferences from this independent study (and an independent set of genomes from our study, Fig. 5), supports the value of our approach and the model we developed. We emphasize the importance of including data from future studies that could measure actual pH preferences of bacterial isolates studied *in vitro*. While curated bacterial phenotypic information from culture collections can provide information on bacterial pH preferences, pre-existing data on bacterial pH preferences of cultured isolates is currently limited to the very narrow distribution of putative pH preferences (85.4% of pH preferences falling between 6 and 8 (*5*)), a pattern that most likely reflects the limited breadth of culturing conditions most commonly used in isolation efforts (*14*). Further quantification of microbial growth responses across large pH gradients under laboratory conditions coupled with whole genome sequencing is key to improving our knowledge of the genetic underpinnings of microbial adaptations to pH.

## Conclusions

We show that biogeographical information can be combined with genomic information to infer and predict the pH preferences of bacterial taxa. This is important given the considerable effort often required to directly measure pH preferences of cultivated taxa *in vitro* and given that such *in vitro* assays are impossible for the majority of bacterial taxa that are resistant to cultivation. We show that pH preferences can be inferred from genomic information alone, making it feasible to leverage the ever-expanding genomic databases, including those which include metagenome-assembled genomes (MAGs) and single cell-assembled genomes (SAGs), to determine a key ecological attribute that cannot be readily determined from taxonomic or phylogenetic information alone (*14*). We not only identified genes that had been previously linked to pH tolerance via detailed studies of select bacteria, but we also identified genes that warrant further investigation as they have not previously been associated with pH adaptations.

Our approach demonstrates the feasibility of using genomic information to make predictions for other important traits that can be difficult to infer directly. In this sense, our work is similar to previous studies that have used genomic data to predict maximum potential growth rates (*65*–*66*), oxygen tolerances (*15*), and temperature optima (*12*), among other traits. However, in those cases, data from cultivated isolates were used to develop the models linking genomic attributes to the trait values of interest. This represents an important bottleneck given that well-characterized cultivated isolates represent only a fraction of the phylogenetic and ecological diversity found in many environments. Instead, we demonstrate how biogeographical information, specifically distributions of taxa across environmental gradients of interest, combined with data from representative genomes, can be used to predict environmental preferences and identify the specific adaptations that may be associated with the ecological attributes. We expect that a similar approach could be used to quantify other relevant traits that have traditionally been difficult to infer directly for uncultivated taxa, including tolerances to changes in moisture, salinity, or heavy metals and other potentially toxic compounds.

The machine learning model presented here has the potential to aid the rational design of microbiomes where information on the pH preferences of bacterial taxa is needed. For example, several studies with N2-fixing rhizobia have successfully improved the symbiotic benefits to legume crops via the isolation and inoculation of acid resistant *Rhizobium* and *Bradyrhizobium* strains (*54*, *55*). Forecasting invasive species spread can also benefit from genome-based predictions of pH (*56*–*58*), given the likely importance of pH as a factor limiting bacterial colonization of new habitats. Also, by predicting bacterial pH preferences, our model can aid the optimization of culturing conditions for any bacterial taxon with available genomic information. The coupling of biogeographical and genomic information can be successfully used to predict the environmental preferences of bacterial taxa, presenting new opportunities for improving our trait-based understanding of microbial life.

## Materials and Methods

### Datasets and sequence processing

We compiled five 16S rRNA gene sequencing datasets that spanned broad gradients in pH and represented distinct ecosystem types. These datasets included two previously published soil datasets (soils collected from across Panama and across Australia, (*18*)), two previously published freshwater datasets (samples collected from streams, rivers, and lakes in Canada (*21*, *22*)), and one new dataset of soils collected from the Savannah River Site (SRS) in South Carolina, United States.

For the SRS dataset, we collected soil samples in May 2021 from patches of savanna and surrounding forests of longleaf and loblolly pine that have been maintained since 2000 as part of the Corridor Project experiment (e.g. (*59*)). The Savannah River Site is a National Environmental Research Park. It is a United States Department of Energy site that is managed by the United States Department of Agriculture Forest Service. Each soil sample consisted of a homogenate of eight 5 cm deep soil core subsamples. We extracted DNA using the DNeasy PowerSoil HTP 96 Kit (Qiagen) from well-mixed soil slurries consisting of 1 g of soil and 2 mL deionized and autoclaved water. We amplified the V4 region of the 16S rRNA gene using universal primers 515 forward and 806 reverse in duplicate PCRs. We cleaned and normalized amplicon samples using SequelPrep Normalization Plate Kit (Applied Biosystems, Waltham, MA, USA) and sequenced a total of 240 soil samples, 22 extraction blanks (400 μL deionized and autoclaved water), and 3 PCR negative controls using paired-end Illumina MiSeq sequencing (300 cycle flow cell). For pH measurements, we created soil slurries consisting of 1 g of soil and 10 mL deionized water, vortexed the slurries at maximum speed for 20 seconds, and then let them rest for about 1 hour.

The five datasets were selected as each one encompassed a broad range in measured sample pH values, each included a sufficiently large number of samples across the five pH gradients (>200 samples per dataset), and pH was strongly correlated with differences in overall bacterial community composition across each dataset (Figure S1). From all but the SRS dataset, we compiled the metadata from open-source databases and from the authors upon request and downloaded the DNA sequences from the Sequence Read Archive (SRA) of the National Center for Biotechnology Information (NCBI). The raw sequences of the SRS project were deposited in the SRA under bioproject ID PRJNA898410. Additional details on these datasets are provided in Figure S1.

All datasets were analyzed using the same bioinformatic pipeline. Briefly, primers, adapters and ambiguous bases were initially removed from the 16S rRNA gene reads using *cutadapt* (v1.18, (*60*)). Sequences were then quality-filtered and amplicon sequence variants (ASVs) were inferred using the DADA2 pipeline (v1.14.1, (*61*)). Chimeric sequences were also removed, and taxonomic affiliations were determined against the SILVA SSU database (release 138, (*62*)). The outputs were loaded into R (*63*) using the *phyloseq* package (v1.38.0, (*64*)) for downstream analyses.

### Statistical inference of pH preferences

Singletons (ASVs represented by only a single read within a given dataset) were removed and samples were rarefied to the minimum read number that ensured sufficient sequencing depth based on rarefaction curves (see Table S1). As archaeal reads were relatively rare in most of these samples, we focused our analyses only on bacteria. Those bacterial ASVs that occurred in fewer than 20 samples per dataset were excluded from the analysis as ASVs needed to occur in a sufficiently large number of samples to effectively infer pH preferences (2-10% of ASVs passed this minimum threshold of occurrence per dataset, Table S1). To infer pH preferences for individual ASVs in each dataset, we first generated 1000 randomized distributions of the relative abundance of each ASV across samples with replacement (i.e. where each relative abundance value could be sampled more than once), and calculated the maximum value for each of these distributions. We then calculated 95% confidence intervals of these relative abundance maxima using the *boot* package (v1.3.28) in R. The extremes of these intervals of relative abundance maxima were then matched to the pH of the samples where these ASVs achieved these relative abundance values, thus obtaining an estimated range of preferential pH for each ASV (i.e. the range of pH in which a given ASV consistently achieved maximal abundance across randomizations). All ASVs with inferred pH preferences that had ranges greater than 0.5 pH units were removed from downstream analyses as we were not confident that we could accurately infer a specific pH preference for those taxa. This stems from the assumption that we could only obtain confident statistical inferences for taxa with narrower pH preferences, excluding taxa with broader ranges in pH preferences to yield more accurate inferences. We visually inspected whether the abundance of each ASV in the non-randomized dataset was within the identified preferred pH range to verify that our pH range filter was sufficiently stringent to exclude unreliable associations between ASVs and pH preferences. The midpoint of the pH range was taken as the estimated pH preference for each ASV. Our preliminary tests showed this bootstrapping approach to be more robust to zero-inflation compared to alternative randomization approaches, GAM, or logistic regression model fits that have previously been used for estimating environmental optima (*36*). We note that many factors, besides just pH, can influence the distributions of bacteria and we restricted our analyses just to those taxa (ASVs) for which pH preferences could effectively be determined (4568 ASVs in total, 468-1614 ASVs per dataset, Table S1).

### Genome search and annotation

The ASVs with estimated pH preferences were matched to the annotated Genome Taxonomy Database (GTDB release 207, (*42*)), using *vsearch* (v2.21.1, (*65*)) to identify representative genomes. We acknowledge that the representative genome identified for any given ASV will not necessarily be an identical match to the genome of that taxon *in situ* as even bacterial taxa with identical 16S rRNA genes can have distinct genomes (*66*). However, this limitation poses a challenge to nearly all genomic analyses of bacteria from environmental samples, as even collections of metagenome-assembled genomes (MAGs) would not necessarily capture the genomic heterogeneity that could exist at finer levels of taxonomic resolution. We also note that, when selecting representative genomes, we only allowed a single base-pair mismatch between the genomes and the 16S rRNA gene amplicons, corresponding to a conservative 99.6% sequence similarity for the 250bp amplicons. In situations where a single ASV matched multiple genomes with the same similarity we selected the most complete genome.

Predicted coding sequences for the ~62,000 unique bacterial genomes (‘species clusters’) available in the GTDB representative set were identified using Prodigal (v2.6.3, (*67*)). The predicted coding sequences for each genome were aligned to the Pfam database (v35.0, (*68*)) using HMMER (v3, (*69*)) to obtain annotation of potential domains, genes, and gene families in each coding sequence. We discarded matches with a bit score lower than 10. This pipeline yielded a list of putative genes and domains found in each of the GTDB genomes that matched the ASVs identified from the samples included in this study with estimated pH preferences (580 genomes in total, 57-293 genomes per dataset, Table S1). The copy numbers of genes and domains were binarized to presence/absence for further analyses.

### Identification of genes associated with pH preference

We next determined associations between the estimated pH preferences of ASVs and the annotated genes in the corresponding representative genomes. We identified genes associated with pH preference by fitting generalized linear models with a binomial distribution and using the logit as a link function with core R functions. For each binarized gene function, we fitted a single model with presence/absence of that gene type as the response variable and the estimated pH preference of each genome as the independent variable. We evaluated the statistical significance of the model coefficients using the Wald test as implemented with R core functions. We obtained the slope (positive or negative) of the relationship between pH preference and the presence/absence of each gene from the model estimates. We verified that the model estimates reliably reflected whether the relationship was positive or negative by plotting the proportion of genomes containing a particular gene type against the estimated pH preferences of those genomes. Finally, we filtered out genes that had non-significant associations to pH preference in all datasets as well as those gene types that had significant associations in only 1 of the 5 datasets (as our goal was to identify consistent associations between genes and pH preferences across multiple datasets). We thus considered those gene types with the same significant association (P < 0.05 and same positive or negative direction of the model estimate) that occurred in 2 or more datasets to be associated with bacterial pH preferences.

### Phylogenetic visualization and analysis

We generated phylogenetic trees for each dataset that included only those ASVs for which pH preferences could be inferred. For each of those ASVs, we selected the corresponding highest quality full-length 16S rRNA gene sequence from the SILVA SSU database (release 138, (*62*)), aligned those sequences with MUSCLE (v5, (*70*)), and created maximum likelihood (ML) trees using the RAxML approach with standard parameters (v8, (*71*)). We visualized and edited the trees using iTOL (v6, (*72*)). We additionally tested whether pH preference had a phylogenetic signal by calculating Blomberg’s K (*73*) and Pagel’s λ (*74*) with the depth at which pH preference had a phylogenetic signal estimated using phylogenetic correlograms (*75*). These three types of phylogenetic analyses were conducted using the R package *phylosignal* (v1.3, (*76*)) using ML trees constructed as described above with all ASVs from a given dataset for which pH preference could be inferred (Table S1).

### Prediction and independent validation of pH preferences from genomic information

We used the full set of genes associated with pH preference across habitats to train a gradient boosted decision tree model to predict bacterial pH preference from genomic information. The gene type presence/absence table was imported into Python and pH preferences were predicted for all ASVs from a one-hot encoded gene matrix using gradient boosted decision trees created with the XGBRegressor function from Python’s *xgboost* package (v1.6.2; (*77*)). Hyperparameter optimization was implemented with the *hyperopt* package (v0.2.7; (*78*)) to select the best hyperparameters for the XGBRegressor model. We validated the accuracy of the model by measuring its mean absolute error (MAE) on an independent 10% of the genomes in each dataset (i.e. the test set). We estimated the impact of each gene type on model predictions using Shapley Additive Explanations (SHAP) as integrated in Python’s *xgboost* package. We also tested the model using a dataset of bacterial taxa with estimated pH preferences from soils across the UK (*40*), from which we selected taxa with unimodal relationships of abundance with pH and therefore more confident estimation of pH preference (models IV and V in (*40*)). For each of these taxa we obtained reference genomes using the same approach described for the ASVs in the other datasets and recorded the presence/absence of the predictive genes in those genomes to run the model.

## Acknowledgments

We thank all the people involved in the acquisition of the data compiled in this study.

## Funding

Swiss National Science Foundation (Early Postdoc Mobility P2EZP3_199849) (JR)

US National Science Foundation (Graduate Research Fellowship Program DGE 1650115) (CCW)

US National Science Foundation (OPP 2133684 and AW5809-826664) (ESO, NF, MH)

NSERC/Hydro-Québec Industrial Research Chair in Carbon Biogeochemistry in Boreal Aquatic Systems (387312-14, IRC 2015-2019) (MS)

NSERC/Hydro-Québec Industrial Research Chair in Carbon Biogeochemistry in Boreal Aquatic Systems (387312-07, IRC 2009-2014) (JPN)

National Science Foundation (awards 1912729 and 1913501) and US Department of Energy to the US Department of Agriculture-Forest Service-Savannah River under Interagency Agreement 89303720SEM000037

## Author contributions

Study design: JR, MH, NF

Data provision: CCW, MS, JPN, NF

Bioinformatics: MH

Machine learning model: ESO

Statistical analyses: JR

Visualization: JR

Writing—original draft: JR, NF

Writing—review & editing: JR, ESO, MH, CCW, MS, JPN, NF

## Data and materials availability

All data are available in the main text or the supplementary materials.

