## Supplementary Materials for "Building a genome-based understanding of bacterial pH preferences"

Josep Ramoneda *et al.*

**This PDF file includes:**

Figs. S1 to S7
Tables S1 to S2

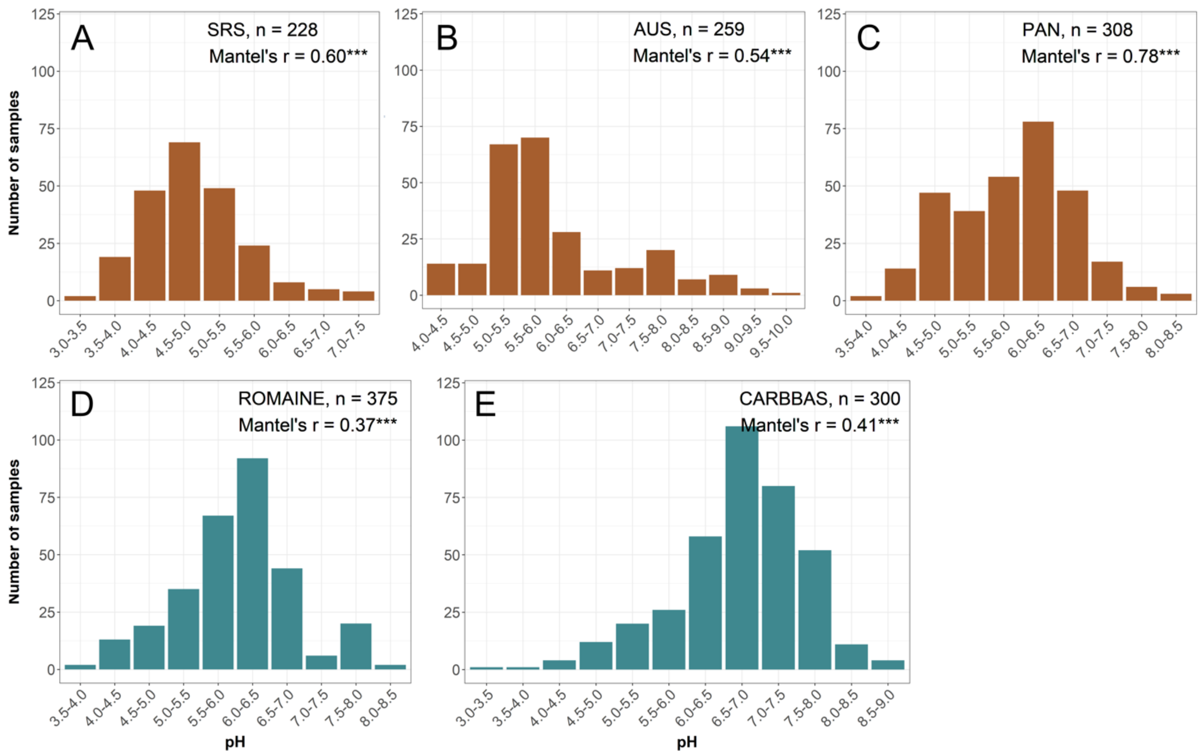


**Fig. S1. Distribution of samples across pH gradients in the datasets used for this study.** The datasets contained 16S rRNA gene sequencing data of bacteria across soils (SRS, USA, *unpublished*; AUS, Australia, (*18*); PAN, Panama, (*18*); depicted in brown), and freshwater (ROMAINE, Canada, (*21)*; CARBBAS; Canada, (*22*); depicted in turquoise). The number of samples (n) included in each dataset is indicated. We also show Mantel tests with 9999 permutations between sample pH and bacterial community composition (pairwise Bray-Curtis distances). In all cases, the Mantel tests were significant (P<0.001), indicating strong associations between sample pH and overall bacterial community composition for each of the 5 datasets.

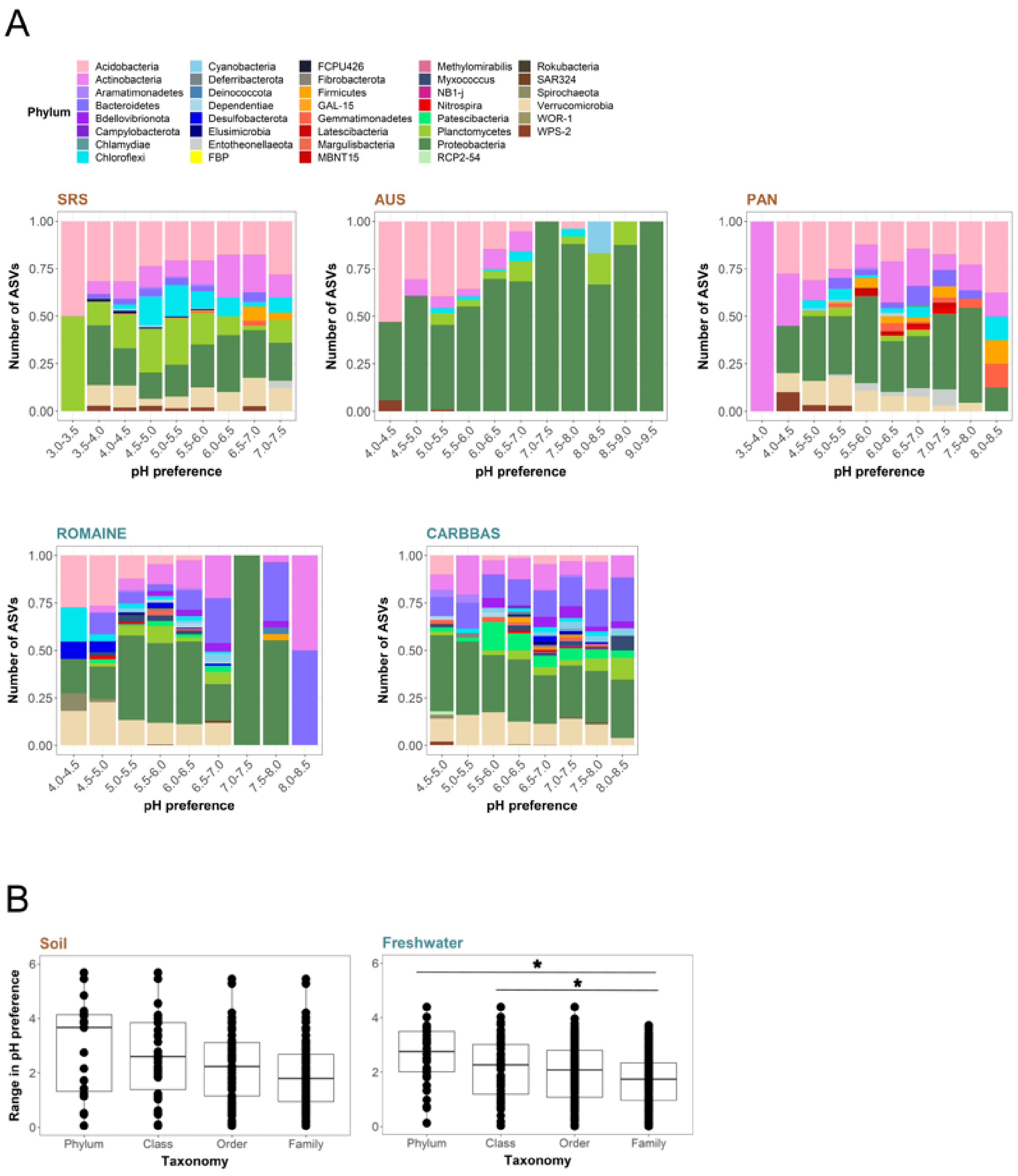


**Fig. S2. Relative abundance of the ASVs with inferred pH preferences classified at the phylum level (A) and range in pH preference of the ASVs across taxonomic levels (B).** The panels in (A) correspond to soil (SRS, AUS, and PAN; in brown) and freshwater (ROMAINE and CARBBAS; in turquoise) datasets. Asterisks in panel (B) indicate a statistically significant difference in the range of pH preference between these broad taxonomic categories based on Welch two-sample t-tests (Holm-Bonferroni-corrected $P < 0.05$).

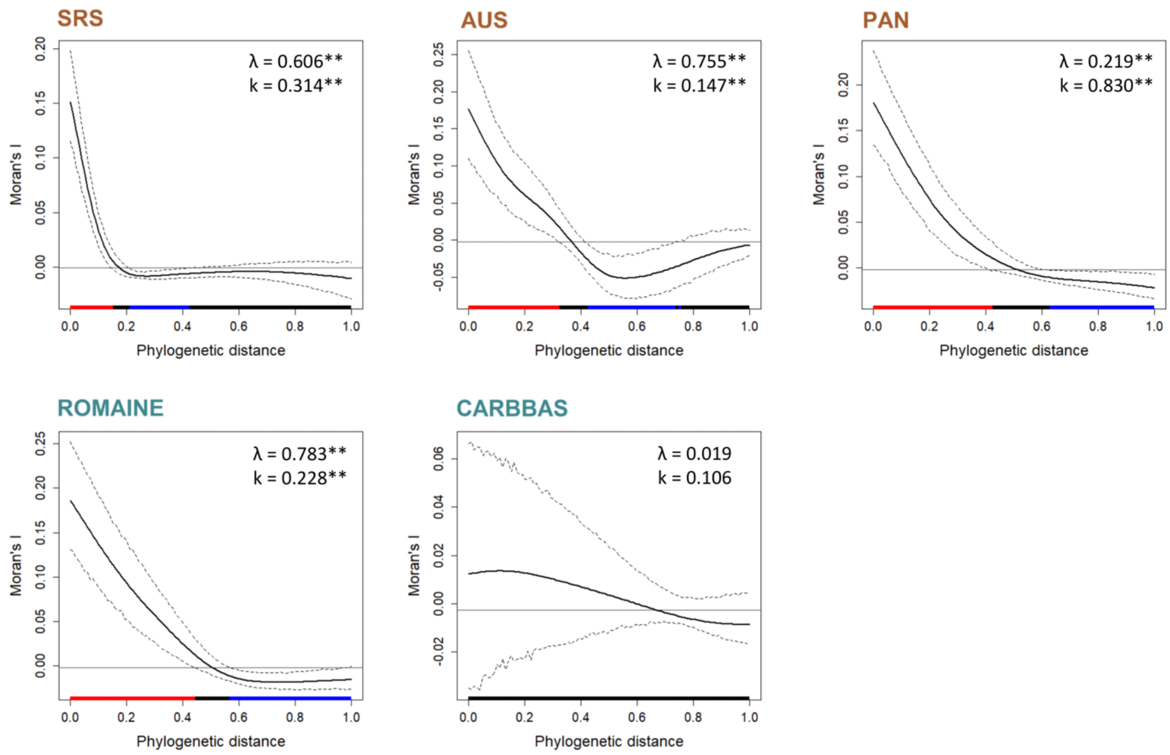


**Fig. S3. Phylogenetic correlograms showing the phylogenetic depth at which pH preference is conserved across soil and freshwater systems.** The text on the graphs shows the Pagel's λ and Blomberg's k statistics for phylogenetic signal in pH preference with statistical significance set at P < 0.05*, P < 0.01**, or P < 0.001***. The mean pairwise phylogenetic distances between taxa within the same family, order, and class are 0.15, 0.25 and 0.30, respectively. Colored lines on the x-axes depict whether pH preference is phylogenetically conserved (red), phylogenetically overdispersed (blue) or showing no phylogenetic signal (black).

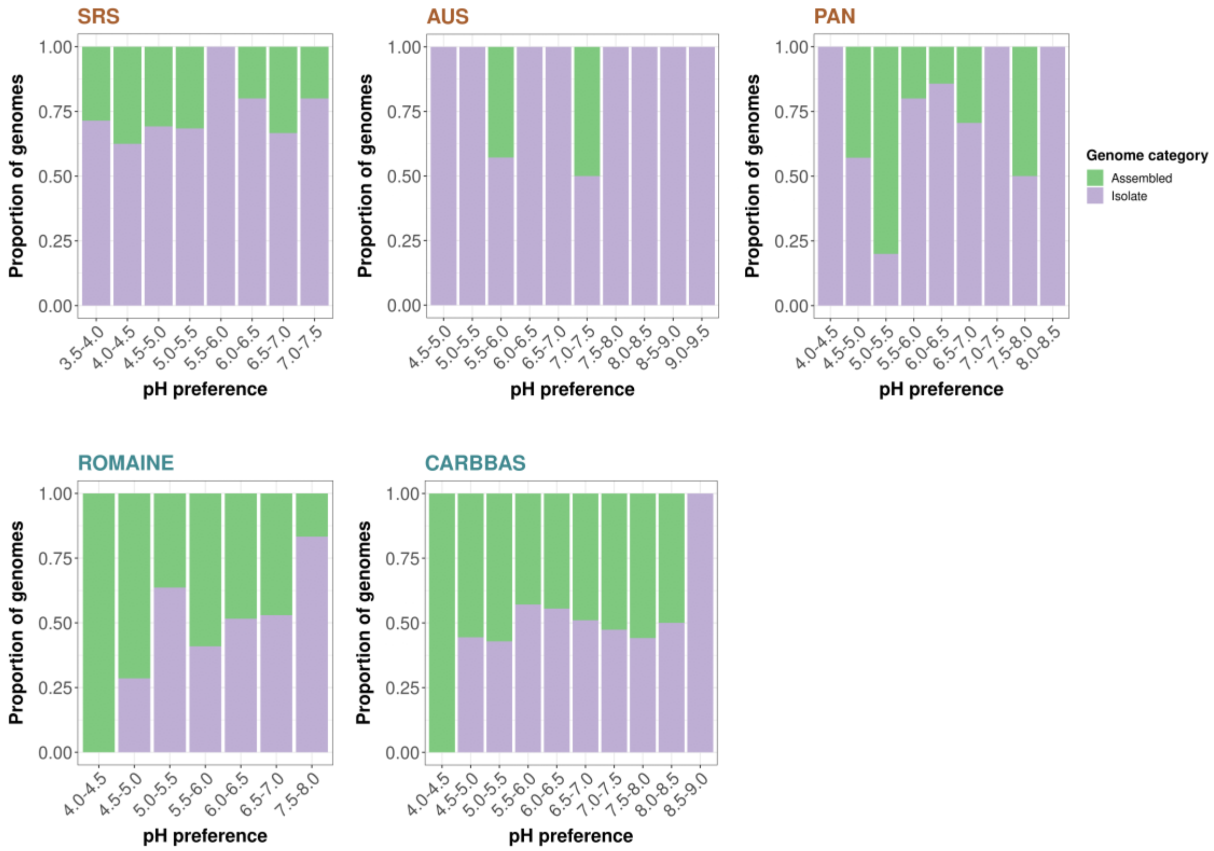


**Fig. S4. Proportion of genomes with inferred pH preferences that were obtained from bacterial isolates ("Isolate") compared to environmental (metagenome) and single cell-assembled genomes ("Assembled") as indicated in the Genome Taxonomy Database (GTDB).**

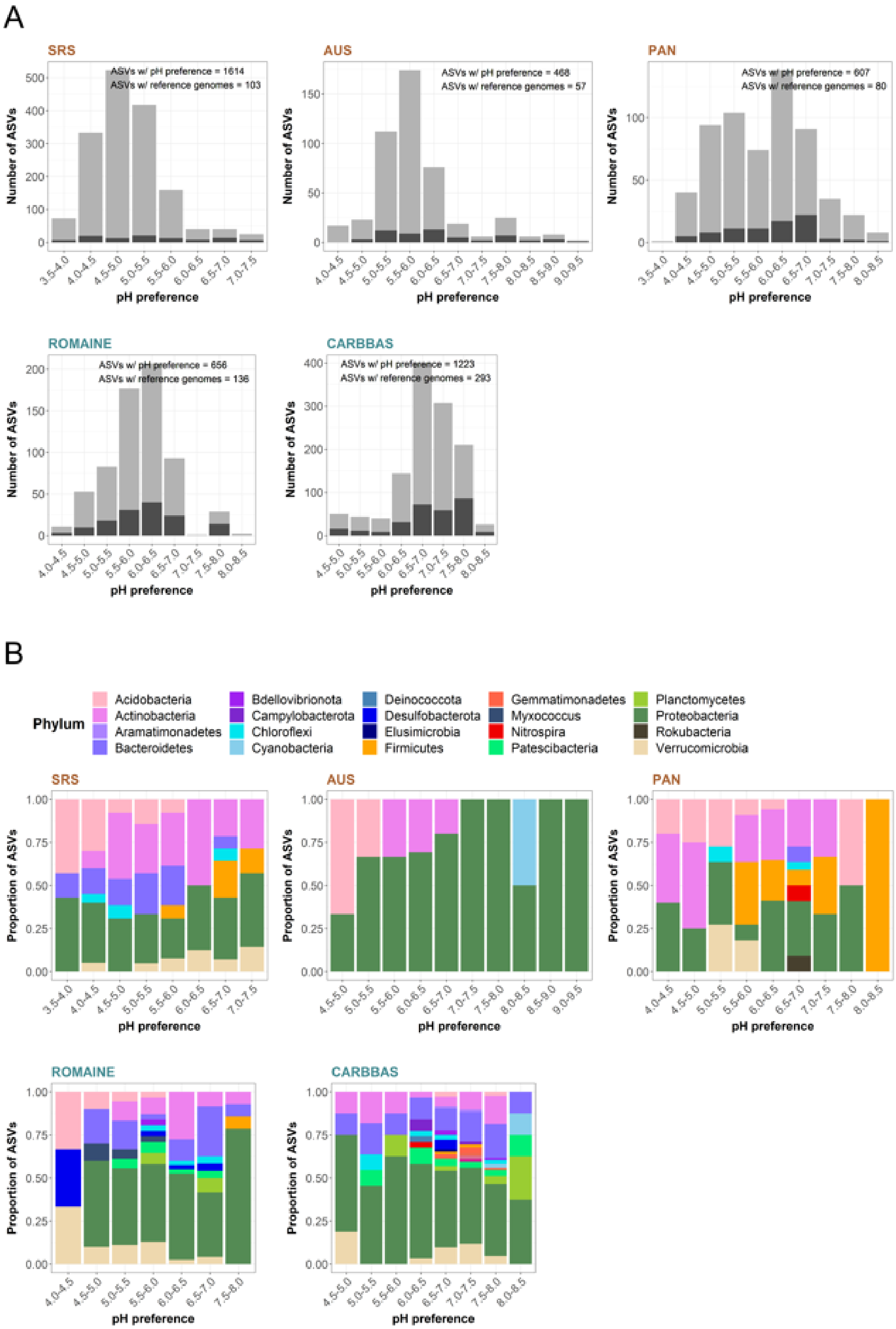


**Fig. S5. The numbers of ASVs with inferred pH preferences (full bars) and those ASVs which also had representative genomes available (dark grey) (A), and the taxonomic affiliations of the representative genomes at the phylum level (B).**

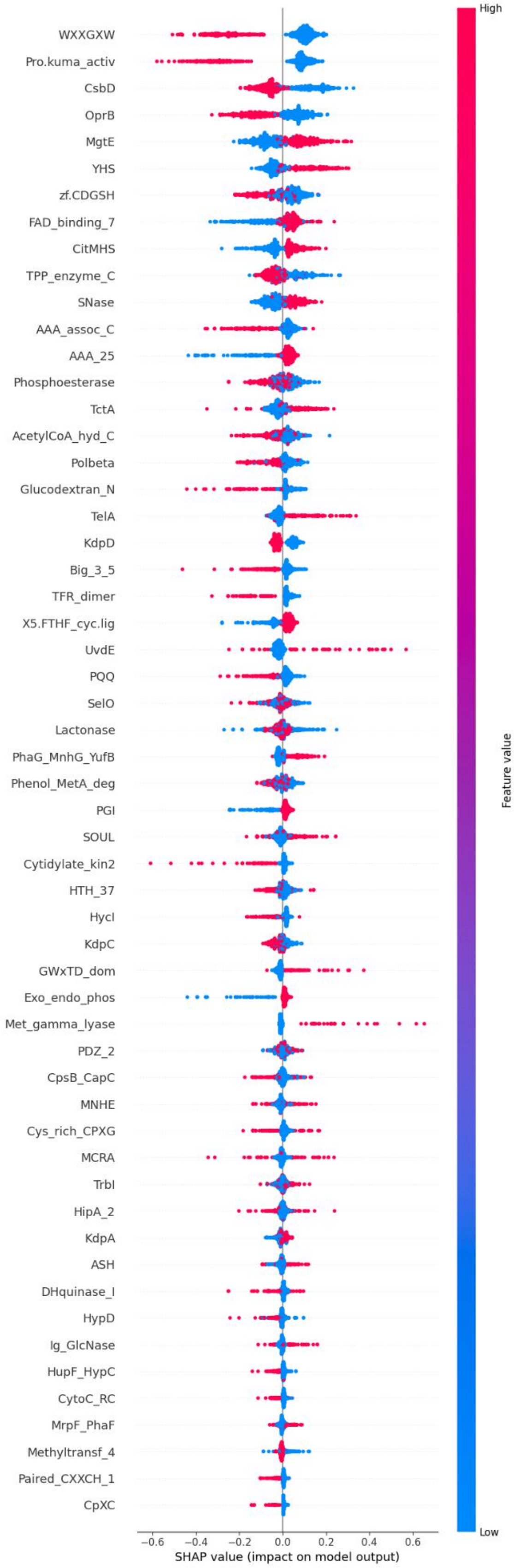


**Fig. S6. Summary of the impact on bacterial pH preference predictions of the 56 genes used to predict bacterial pH preferences with machine learning.** Positive values have a positive effect on pH preference predictions (higher inferred pH preferences), and negative values have a negative effect on pH preference predictions (lower inferred pH preferences). The coloring indicates the impact on model predictions of the presence (red) and absence (blue) of a given gene type in a genome.

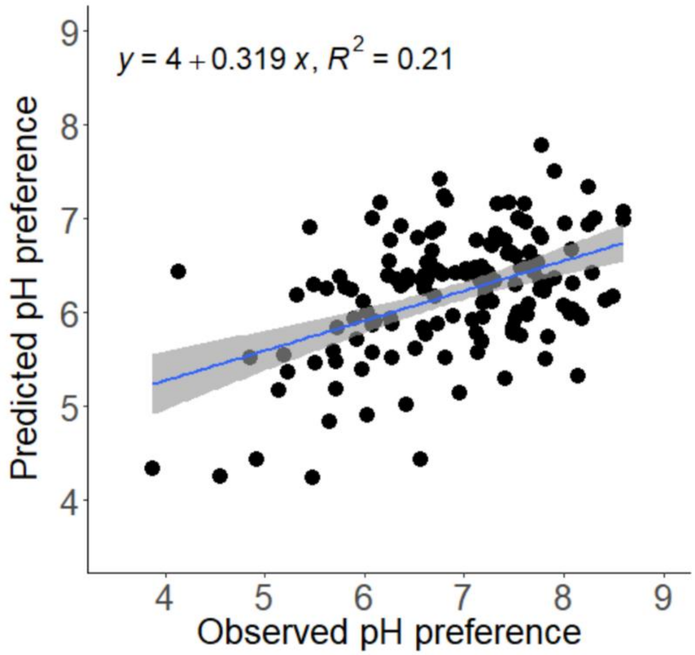


**Fig. S7. Correlation between the estimated and predicted pH preferences of bacterial taxa based on a gradient boosted decision tree model based on the presence/absence of 56 top predictor gene types.** These data are for the 148 representative genomes obtained for taxa with estimated pH preferences in a study of soils across the UK (Jones *et al.* 2021).

| Dataset | Rarefaction depth (# reads per sample) | #ASVs after rarefaction | #ASVs after ubiquity threshold | #ASVs with inferred pH preferences | #ASVs for which pH preferences could be inferred with matching genomes |
|---|---|---|---|---|---|
| SRS (USA) | 14973 | 32865 | 3159 | 1614 | 103 |
| AUS (Australia) | 4420 | 19219 | 864 | 468 | 57 |
| PAN (Panama) | 7834 | 18185 | 944 | 607 | 80 |
| ROMAINE (Canada) | 13254 | 44688 | 1295 | 656 | 136 |
| CARBBAS (Canada) | 31779 | 135318 | 2713 | 1223 | 293 |

**Table S1. Features of the ASVs included in the study.** The ubiquity threshold forced ASVs to occur in at least 20 samples for each study.

| Gene type | Function | Category | Association to pH preference | Pfam ID | Available literature |
|---|---|---|---|---|---|
| 5-FTHF_cyc-lig | Folate metabolism | Folate metabolism | (+) | PF01812 | - |
| AAA_25 | ATPases | ATPases | (+) | PF13481 | (*79*, *80*) |
| AAA_assoc_C | ATPases | ATPases | (-) | PF09821 | (*79*, *80*) |
| AcetylCoA_hyd_C | Acetyl-CoA hydrolase/transferase | Acetyl-CoA metabolism | (-) | PF13336 | (*28*) |
| ASH | Ciliary/Flagellar function | Motility | (-) | PF15780 | - |
| Big_3_5 | Ig like domains | Surface proteins | (-) | PF16640 | - |
| CitMHS | Citrate transporter | Transmembrane anion transporter | (+) | PF03600 | (*28*, *81*) |
| CpsB_CapC | Phosphatases in polysaccharide synthesis | Phosphatases | (-) | PF19567 | (*28*) |
| CpXC | Unknown | Unknown | (-) | PF14353 | - |
| CsbD | Unknown | Stress response | (-) | PF05532 | (*82*) |
| Cys_rich_CPXG | Unknown | Unknown | (-) | PF14255 | - |
| Cytidylate_kin2 | Cytidylate kinase | Kinases | (-) | PF13189 | - |
| CytoC_RC | Cytochrom C photosynthetic reaction center | Cytochrom C | (-) | PF02276 | - |
| DHquinase_I | Dehydrokinase | Kinases | (-) | PF01487 | (*28*) |
| Exo_endo_phos | Endonuclease/Exonuclease/phosphatase family | Phosphatases | (+) | PF03372 | (*28*) |
| FAD_binding_7 | Photolyase | Nucleases | (+) | PF03441 | (*28*) |
| Glucodextran_N | Hydrolases of dextrans | Sugar metabolism | (-) | PF09137 | (*83*) |
| GWxTD_dom | Unknown | Unknown | (-) | PF20094 | - |
| HipA_2 | Kinase that inhibits tRNA synthase (antibiosis) | Kinases | (-) | PF20613 | (*28*) |
| HTH_37 | Helix-turn-helix domain | Unknown | (-) | PF13744 | - |
| HupF_HypC | Hydrogenase expression/formation and maturation proteins | Hydrogenase maturation | (-) | PF01455 | (*28*, *84*) |
| HycI | Hydrogenase maturation protease | Hydrogenase maturation | (-) | PF01750 | (*84*) |
| HypD | Hydrogenase formation hypA family | Hydrogenase maturation | (-) | PF01924 | (*28*, *84*) |
| Ig_GlcNase | Glucosaminidase | Hydrolases | (-) | PF18368 | (*85*) |
| KdpA | $K^+$ transporter | Transmembrane cation transporters | (-) | PF03814 | (*86*–*88*) |
| KdpC | $K^+$ transporter | Transmembrane cation transporters | (-) | PF02669 | (*86*–*88*) |
| KdpD | $K^+$ transporter | Transmembrane cation transporters | (-) | PF02702 | (*86*–*88*) |
| Lactonase | Acylated homoserine lactone metabolism | Hydrolases | (-) | PF10282 | (*28*) |
| MCRA | Lipid hydratase | Antigens | (-) | PF06100 | - |
| Met_gamma_lyase | Lyase of methionine metabolism | Methionine metabolism | (+) | PF06838 | - |
| Methyltransf_4 | Methyltransferase | Methyltransferases | (+) | PF02390 | - |
| MgtE | Transmembrane $Mg^{+2}$ transporter | Transmembrane cation transporters | (+) | PF01769 | - |
| MNHE | $Na^+/H^+$ antiporter | $Na^+/H^+$ antiporters | (+) | PF01899 | (*28*, *89*) |
| MrpF_PhaF | $Na^+/H^+$ antiporter | $Na^+/H^+$ antiporters | (+) | PF04066 | (*33*, *90*, *91*) |
| OprB | Carbohydrate porin | Carbohydrate transport | (-) | PF04966 | - |

| | | | | | |
|---|---|---|---|---|---|
| Paired_CXXCH_1 | Motif part of C type cytochrome | Cytochrom C | (-) | PF09699 | - |
| PDZ_2 | Transmembrane proteins | Transmembrane proteins | (-) | PF13180 | (*28*) |
| PGI | Gluc to Fruc 6-phosphate | Sugar metabolism | (+) | PF00342 | (*92*) |
| PhaG_MnhG_YufB | $Na^+/H^+$ antiporter | $Na^+/H^+$ antiporters | (+) | PF03334 | (*28*, *90*) |
| Phenol_MetA_deg | Phenol degradation | Phenol degradation | (-) | PF13557 | - |
| Phosphoesterase | Phospholipase C | Phosphatases | (-) | PF04185 | (*93*, *94*) |
| Polbeta | Nucleotidyltransferase | Nucleotidyltransferase | (-) | PF18765 | - |
| PQQ | Beta propeller domain of quinolones | Quinolones | (-) | PF01011 | (*28*) |
| Pro-kuma_activ | Serine protease | Proteases | (-) | PF09286 | (*95*) |
| SelO | Kinase that does AMPylation to proteins | Kinases | (+) | PF02696 | - |
| SNase | Ca-dependent nuclease | Nucleases | (+) | PF00565 | - |
| SOUL | Heme binding proteins | Heme binding proteins | (+) | PF04832 | - |
| TctA | Transmembrane citrate transporter | Transmembrane anion transporter | (+) | PF01970 | (*28*) |
| TelA | Toxic anion resistance (tellurite) | Transmembrane anion transporter | (+) | PF05816 | - |
| TFR_dimer | Transferrin dimerisation domain | Fe receptors | (-) | PF04253 | (*96*) |
| TPP_enzyme_C | Thyamin pirophsophate binding domain | Phosphate receptors | (-) | PF02775 | - |
| TrbI | Conjugation-related protein | Type IV secretion system | (-) | PF03743 | - |
| UvdE | UV damage repair endonuclease | Nucleases | (+) | PF03851 | - |
| WXXGXW | Unknown | Unknown | (-) | PF12779 | - |
| YHS | $Cu^{+2}$ transporter | Transmembrane cation transporters | (+) | PF04945 | (*28*) |
| zf-CDGSH | Fe-binding membrane protein acting as a redox-active pH-labile 2Fe-2S cluster | Fe receptors | (-) | PF09360 | - |

**Table S2. List of genes significantly associated with bacterial pH preference across habitats.** When known, the main functions associated with the genes are reported, and the direction of their relationship to pH preference is indicated as either positive (+) or negative (-). When available, we also included the relevant references reporting a putative role of the genes in bacterial pH preference. These 56 genes were included in a regression predictive modelling framework based on gradient boosted decision trees that predicts pH preference of bacterial taxa based on presence/absence information on those genes.
